# Functional connectivity between the amygdala and prefrontal cortex underlies processing of emotion ambiguity

**DOI:** 10.1101/2023.01.24.525116

**Authors:** Sai Sun, Hongbo Yu, Rongjun Yu, Shuo Wang

## Abstract

Processing facial expressions of emotion draws on a distributed brain network. In particular, judging ambiguous facial emotions involves coordination between multiple brain areas. Here, we applied multimodal functional connectivity analysis to achieve network-level understanding of the neural mechanisms underlying perceptual ambiguity in facial expressions. We found directional effective connectivity between the amygdala, dorsomedial prefrontal cortex (dmPFC), and ventromedial PFC, supporting both bottom-up affective processes for ambiguity representation/perception and top-down cognitive processes for ambiguity resolution/decision. Direct recordings from the human neurosurgical patients showed that the responses of amygdala and dmPFC neurons were modulated by the level of emotion ambiguity, and amygdala neurons responded earlier than dmPFC neurons, reflecting the bottom-up process for ambiguity processing. We further found parietal-frontal coherence and delta-alpha cross-frequency coupling involved in encoding emotion ambiguity. We replicated the EEG coherence result using independent experiments and further showed modulation of the coherence. EEG source connectivity revealed that the dmPFC top-down regulated the activities in other brain regions. Lastly, we showed altered behavioral responses in neuropsychiatric patients who may have dysfunctions in amygdala-PFC functional connectivity. Together, using multimodal experimental and analytical approaches, we have delineated a neural network that underlies processing of emotion ambiguity.

**Significance Statement:** A large number of different brain regions participate in emotion processing. However, it remains elusive how these brain regions interact and coordinate with each other and collectively encode emotions, especially when the task requires orchestration between different brain areas. In this study, we employed multimodal approaches that well complemented each other to comprehensively study the neural mechanisms of emotion ambiguity. Our results provided a systematic understanding of the amygdala-PFC network underlying emotion ambiguity with fMRI-based connectivity, EEG coordination of cortical regions, synchronization of brain rhythms, directed information flow of the source signals, and latency of single-neuron responses. Our results further shed light on neuropsychiatric patients who have abnormal amygdala-PFC connectivity.

## Introduction

Faces are among the most important visual stimuli that we perceive in everyday life. We are able to not only perceive subtle facial expressions but also recognize conflicting and ambiguous facial expressions. The processing of faces and facial emotions engages a distributed network of brain regions [1–3]. In particular, judging ambiguous facial expressions requires orchestration between multiple brain areas, notably involving the amygdala and two regions of the prefrontal cortex (PFC): the dorsomedial PFC (dmPFC) and ventromedial PFC (vmPFC) [4, 5].

The amygdala has long been associated with a key role in recognizing facial emotions [2, 6, 7]. Human studies demonstrated a selective impairment in recognizing fearful faces in participants that lack a functional amygdala [8], mirrored by neuroimaging studies showing significant activation differences within the amygdala to fearful faces compared to happy faces [9]. Neurons in the human amygdala encode subjective judgment of facial emotions rather than simply their stimulus features [10]. Consistent with human single-unit studies, intracranial field potentials in the human amygdala show modulation by emotion and attention [11], and neurons in the monkey amygdala encode facial expressions [12, 13]. In addition to facial expressions, the amygdala is crucial for identifying ambiguous stimuli and in modulating vigilance and attention as a function thereof [14–16]. For instance, the BOLD-fMRI signal in the amygdala is correlated with the level of ambiguity in decision making [17] and focal amygdala damage undermines decision making under ambiguity [18]. In particular, our previous work has shown that both neurons and BOLD-fMRI from the human amygdala parametrically encode the intensity of specific facial emotions and their categorical ambiguity [5]. Furthermore, unpredictability of stimuli even without any motivational information activates the basolateral amygdala in mice and causes sustained neural activity in the amygdala in humans [19]. Together, these findings suggest that the amygdala plays a key role in processing ambiguity.

The dmPFC, notably including the dorsal anterior cingulate cortex (dACC) and pre-supplementary motor area (pre-SMA), plays a critical role in cognitive control, including the detection of performance errors and the monitoring of conflict [20–24], reward-based decision making and learning more generally [25], as well as emotion processing and regulation [26]. The vmPFC plays a multifaceted role in emotion, decision making, and social cognition [27]. It is involved in fear extinction [28], value comparison and confidence [29], as well as emotion regulation [26]. Human neurological patients with a focal vmPFC damage demonstrate a severe defect in decision making with ambiguity [30]. While a single unifying principle of dmPFC and vmPFC function remains elusive, most of the above functions involve the processing of ambiguity in some form. Ambiguity inherently involves conflict in how sensory information maps onto categories or choices, requires continuous monitoring of ongoing actions, and triggers dynamic adjustments in cognitive control. Ambiguous emotional faces relative to unambiguous emotional faces activate the dmPFC, whereas ambiguous affective decisions relative to ambiguous gender decisions activate the vmPFC [31]. In particular, we have shown that BOLD-fMRI from both dmPFC and vmPFC encode emotion ambiguity [4], and a neural signature originating from the dmPFC and vmPFC, the late positive potential (LPP), indexes decision ambiguity of facial expressions of emotion [4, 32].

While our prior studies have revealed compelling functional localization of emotion ambiguity [4, 5, 32], it remains largely unclear how these distributed brain areas interact and coordinate with each other to collectively encode emotion ambiguity, especially when the task requires orchestration between multiple brain areas. The behavioral findings from human patients with a focal damage in the bilateral amygdala [5] further suggested that a network view is needed to understand the underlying neural processes. The functional network including the dmPFC, vmPFC, and the amygdala is critical for emotion processing, which has been supported by the pattern of anatomical connectivity [28]. Robust reciprocal connections have also been found between the dmPFC and the lateral basal nucleus of the amygdala, and functional connectivity data from humans show a similar pattern [22]. The vmPFC is important for the generation and regulation of negative emotion, through its interactions with the amygdala [27, 33]. Functional neuroimaging has shown that the activity in the amygdala, vmPFC, and dmPFC reflects the amount of emotional conflict, and the vmPFC modulates the activity in the amygdala to resolve such conflict [34]. Furthermore, effective amygdala-PFC (including both dmPFC and vmPFC) connectivity predicts individual differences in successful emotion regulation [35]. However, a detailed network-level understanding remains missing.

To fill this gap, in this study, we employed multimodal experimental approaches, including fMRI, EEG, and human single-neuron recordings, to comprehensively investigate the functional network underlying emotion ambiguity. We sought to identify the functional/effective connectivity of the amygdala-PFC network in representing and resolving ambiguous facial expressions in neurotypical individuals, and we hypothesize that the amygdala is functionally connected with the vmPFC and dmPFC when modulated by levels of emotion ambiguity. We further examined behavioral performance in several groups of neuropsychiatric patients who show dysfunction of the amygdala-PFC network. Specifically, task-based and task-free neuroimaging studies have revealed altered amygdala-PFC functional or effective connectivity in people with autism spectrum disorder (ASD) [36, 37], attention-deficit/hyperactivity disorder (ADHD) [38], social anxiety disorder [39, 40], major depression [41–43], aggressive behavior [44], schizophrenia [45], and post-traumatic stress disorders (PTSD) [46, 47]. Therefore, we hypothesize that patients with a dysfunctional amygdala-PFC network show altered behavioral response to ambiguous facial expressions.

## Methods

### Participants

In the main task (face judgment task with fear-happy morphed emotions), 19 neurotypical participants (4 male, 20.9±2.02 [mean±SD] years) participated in the functional magnetic resonance imaging (fMRI) experiment, 16 neurosurgical patients (11 male, 42.3±17.0 years; 22 sessions) participated in the single-neuron recording experiment, and 23 neurotypical participants (6 male, 22.4±2.17 years) participated in the electroencephalogram (EEG) experiment. Furthermore, 16 neurotypical participants (5 male, 19.63 ± 0.96 years) performed the EEG control experiment with a speeded response as well as the EEG control experiments with different task instructions (i.e., judging the gender or the wealth [rich versus poor] of the face model). Lastly, 32 neurotypical participants (17 male, 20.6±1.79 years) performed the EEG control experiment with context modulation.

Three groups of neuropsychiatric patients (autism spectrum disorder [ASD], in-patient schizophrenia [SCZ], and out-patient SCZ) and one control group of neurotypicals participated in the in-lab experiment. Specifically, 18 high-functioning participants with ASD (15 male, 30.8±7.40 years; ASD diagnosis confirmed by both DSM-V/ICD-10 and Autism Diagnostic Observation Schedule-2 [ADOS-2]), 29 in-patient SCZ participants, 24 out-patient SCZ participants, and 15 neurotypical controls (35.1±11.4 years) performed the main task. We excluded 7 in-patient SCZ participants and 3 out-patient SCZ participants from further analysis because of their misunderstanding of instructions, repeated button presses, and incomplete data records. Therefore, data from 22 in-patient participants (12 male, 36.1±10.48 years) and 21 out-patient participants (10 male, 36.7±8.14 years) were further analyzed. There was no significant difference in age between SCZ groups (two-tailed two-sample *t*-test, *t*(41) = 0.366, P = 0.718, *d* = 0.07). All participants with SCZ were diagnosed by clinical psychiatrists and had semi-structured clinical interviews with psychotherapists in the hospital. In-patient SCZ participants had a history of schizophrenia of 5 to 31 years (mean±SD: 14.0±7.63 years). Furthermore, there was no significant difference in age (one-way ANOVA, *F*(3, 75) = 1.19, P = 0.323, η_p_^2^ = 0.14) across in-lab neuropsychiatric patient populations and neurotypicals.

In addition to in-lab participants, four groups of self-identified neuropsychiatric patients (anxiety, depression, ASD, and attention-deficit/hyperactivity disorder [ADHD]) and one control group neurotypicals were recruited online using the Prolific platform (https://www.prolific.co/). The experiments were programmed using Labvanced (https://www.labvanced.com/), which offers a graphical task builder with high temporal accuracy or response time measures. Specifically, 38 participants with self-reported anxiety (16 male, 37.31±6.62 years), 35 participants with self-reported depression (19 male, 36.17±7.75 years), 34 participants with self-reported autism (16 male, 36.76±6.81 years), 36 participants with self-reported ADHD (21 male, 34.80±6.45 years), and 56 control participants without any self-reported neuropsychiatric disorders (25 male, 36.96±6.67 years) performed the online version of the main task. All participants were proficient in English, unique, and involved in only one experiment. There was no significant difference in age (one-way ANOVA, *F*(4, 198) = 0.781, P = 0.539, η_p_^2^ = 0.05) across groups.

All participants had normal or corrected-to-normal visual acuity. Participants provided written informed consent according to protocols approved by the Institutional Review Board (IRB) of the South China Normal University, Cedars-Sinai Medical Center, California Institute of Technology, Fujian University of Traditional Chinese Medicine, and Tohoku University.

### Stimuli and task

Stimuli were morphed expression continua between exemplars of fearful and happy expressions. Four individuals (two female) were chosen from the STOIC database [48], a database of face images expressing highly recognizable emotions. The facial expressions were morphed by a computer algorithm, therefore, they are not real faces. For each individual we selected unambiguous exemplars of fearful and happy expressions as evaluated with normative rating data provided by the database creators. To generate the morphed expression continua for this experiment, we interpolated pixel value and location between fearful exemplar faces and happy exemplar faces using a piece-wise cubic-spline transformation over a Delaunay tessellation of manually selected control points. We created 5 levels of fear-happy morphs, ranging from 30% fear/70% happy to 70% fear/30% happy in steps of 10%. Low-level image properties were equalized by the SHINE toolbox [49] (The toolbox features functions for specifying the (rotational average of the) Fourier amplitude spectra, for normalizing and scaling mean luminance and contrast, and for exact histogram specification optimized for perceptual visual quality).

In the main task, on each trial, a face was presented for 1 second followed by a question prompt asking participants to make the best guess of the facial emotion. Participants reported faces as fearful or happy by pressing a button on the keyboard or response box. After stimulus offset, participants had 2 seconds (for in-lab participants) or 5 seconds (for online participants) to respond, otherwise, the trial would be aborted and discarded. Participants were instructed to respond only after stimulus offset. Patients with schizophrenia did not have a time constraint to respond. No feedback message was displayed, and the order of faces was completely randomized for each participant. After judging the emotions, participants were asked to indicate their confidence of judgment by pushing the button 1 for “very sure”, 2 for “sure”, or 3 for “unsure”. As with the emotion judgment, participants had 2 seconds to respond before the trial was aborted, and no feedback message was displayed. Confidence rating was omitted for EEG participants, fMRI participants, 7 in-lab ASD participants, and 4 in-lab neurotypical participants. An inter-trial-interval (ITI) was jittered randomly with a uniform distribution between 1 and 2 seconds for all participants except 2-8 seconds for fMRI participants. EEG participants performed 252 trials in 2 blocks, fMRI participants performed 168 trials in 2 blocks, neurosurgical patients performed 176 to 440 trials in 2 to 5 blocks, in-lab ASD and neurotypical participants performed 252 trials in 3 blocks, SCZ participants performed 84 trials in 2 blocks, and online participants performed 84 trials in 1 block. All trials were pooled for analysis.

In the speeded version of the task, participants were instructed to respond as quickly as possible. The stimulus stayed on the screen until button press. Similarly, participants had 2 seconds to respond, otherwise the trial was aborted and discarded. In contrast to the main task, the question prompt asking participants to make the best guess of the facial emotion preceded the stimulus and was presented for 500 ms. Participants performed 280 trials in 2 blocks.

In the context modulation task, participants only judged anchor faces in the first and third block (64 trials each), whereas they judged both anchor faces and morphed faces (identical to the main task) in the second block (192 trials).

In the gender judgment task, participants were asked to judge the gender of the face. This task had no ambiguity because all four face models had clearly recognizable genders. In contrast, participants were asked to guess the wealth (poor vs. rich) of the face model in the wealth judgment task, and this task had the highest ambiguity because whether the face model is poor or rich could not be told without any priors. Here, we used the speeded version of the task. There were 280 trials in 2 blocks for each experiment.

### Behavioral analysis

We are mainly interested in the inter-individual *threshold* in emotion discrimination and *sensitivity* to the changes in emotion intensity. We fitted a logistic function to obtain smooth psychometric curves and derived the two parameters from the fitted curves:

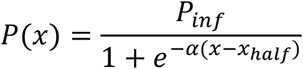

where *P* is the percentage of trials judging faces as fear, *x* is the morph level, *P_inf_* is the value when *x* approaches infinity (the curve’s maximum value), *x_half_* is the symmetric inflection point (the curve’s midpoint, *threshold*), and α is the steepness (*sensitivity*) of the curve. *P_inf_*, *x_half_*, and *α* were fitted from the observed data (*P* and *x*). Smaller *x_half_* suggests that participants were more likely to judge faces as fearful (i.e., a lower threshold to report fearful), and vice versa for larger *x_half_*. Flatter curves (smaller α) suggest that participants were less sensitive to the change in emotion intensity since they made similar judgments given different morph levels, and vice versa for steeper curves (larger α). We derived these two parameters (threshold and sensitivity) for each participant.

### Functional magnetic resonance imaging (fMRI)

Detailed methods of fMRI imaging acquisition and functional localization have been described in our previous reports [4, 5]. Below, we describe the methods for functional connectivity analyses, which have not been reported in our previous studies.

Our previous work [4] revealed a significant increase in BOLD signal in the bilateral inferior frontal gyrus (IFG)/anterior insula and dorsal medial prefrontal cortex (dmPFC) as a function of *increasing* emotion ambiguity; and we also found a significant increase in BOLD signal in the right amygdala, left vmPFC, posterior cingulate cortex (PCC), dorsolateral prefrontal cortex (dlPFC), and inferior parietal lobule (IPL) as a function of *decreasing* emotion ambiguity. Moreover, we observed an increase in BOLD signal in the left amygdala, dmPFC, and insula as a function of decreasing fearful intensity. In this study, our functional connectivity analyses mainly focused on the amygdala-PFC network that has been identified in the main contrast of emotion ambiguity.

### fMRI: psychophysiological interaction (PPI) analyses

A psychological context does not only modulate the strength of regional brain activation, but can also modulate the physiological connectivity between two brain regions. The physiological connectivity between two brain regions that vary with the psychological context is known as psychophysiological interaction (PPI) [50]. In this study, the regional connectivity arisen from perceiving emotional faces could be modulated by the levels of emotion ambiguity. Thus, we conducted a PPI analysis to identify the “target” regions whose connectivity with a seed region (i.e., the right amygdala) varied as a function of increasing/decreasing emotion ambiguity.

A GLM was constructed with PPI regressors of (1) the main physiological effect of the right amygdala for the main contrast of decreasing ambiguity levels, (2) the main psychological effect on decreasing ambiguity, and (3) the interaction effect between the right amygdala and target regions, corresponding to *PPI.Y*, *PPI.P*, and *PPI.ppi* in the design matrix, respectively. The first-level model included the main effect of the physiological, psychological, and interactive effects convolved by the hemodynamic response function (HRF), as well as six movement parameters as effects of no interest. Participant-specific PPI contrast images were computed and entered into a second-level GLM to identify brain areas for which the change in connectivity with the right amygdala was modulated by emotion ambiguity. The same statistical approaches previously described for the functional localization analyses [4] were employed for the second-level connectivity maps. Specifically, activations were reported if they survived P < 0.001 uncorrected, cluster size *k* > 20, or P < 0.05 FWE after small volume correction (SVC). The pre-defined regions of interest (ROIs) in the dmPFC and vmPFC for SVC were chosen based on previous studies [31, 51].

A similar PPI analysis was conducted for increasing/decreasing fear intensity with the left amygdala (**Supplementary Fig. 1**).

### fMRI: high-order PPI analyses

To identify the regions that had functional connectivity with the right amygdala and whose connectivity was modulated by the inter-individual difference in ambiguity sensitivity, high-order PPI model was constructed. Ambiguity sensitivity of each participant was defined as the difference in RT between high ambiguity and anchor, and it was used in a regression analysis to identify the brain regions that responded to the main PPI contrast (i.e., PPI.ppi generated from the primary PPI model). Such subsequent PPI model was referred to as “high-order” because it was built on the main PPI contrast.

### fMRI: dynamic causal modeling (DCM) analyses

DCM explains the activity of groups of regions in terms of (1) “driving” inputs (here, processing a face, regardless of emotion ambiguity) directly triggering the response in one or more areas of the network, and (2) a psychological context (here, levels of emotion ambiguity) acting on “intrinsic” pathways and further modulating the pattern of effective connectivity between regions. Three matrices were built with Matrix A representing the intrinsic coupling between regions, Matrix B representing the changes in functional coupling due to psychological context, and Matrix C representing the direct influences of “driving” inputs on the network under a specific psychological context.

To establish the model, we extracted data from the right amygdala within a 10 mm sphere centered on the local maximum of regional activation under the decreasing ambiguity contrast (peak: Montreal Neurological Institute [MNI] coordinate: *x* = 30, *y* = 0, *z* = –21). We also extracted the volumes of interest (VOIs) from the left vmPFC under the decreasing ambiguity contrast (peak: *x* = –6, *y* = 39, *z* = –9) and the left dmPFC under the increasing ambiguity contrast (peak: *x* = –6, *y* = 15, *z* = 54) within a 10 mm sphere centered on the local maxima of regional activation. It is worth noting that similar results could be derived using the right vmPFC (peak: *x* = 21, *y* = 48, *z* = –3) and dmPFC (peak: *x* = 12, *y* = 36, *z* = 48) indicated by the PPI results (**Supplementary Fig. 3**). The standard model included “intrinsic” bidirectional connections among the amygdala, vmPFC, and dmFPC. These “intrinsic” connections (DCM matrix A) represent the intrinsic coupling between regions in the absence of any experimental manipulations. Beyond the intrinsic connections, effective connectivity in a network can be changed in two ways. First, the “driving inputs” (faces vs. fixation, DCM matrix C) can directly influence an individual region or a group of regions within the network. Second, changes due to psychological context (i.e., increasing/decreasing ambiguity levels; DCM matrix B) can modulate both “intrinsic” and functional connections within the network.

We specified 44 models in which the number and direction of involved regions systematically varied. To identify the best family of model fits, we conducted a random-effect family inference analysis. We fixed the driving inputs to the network (i.e., face signals) through the amygdala node. Based on the inputs to the amygdala, we had four families of models (**Supplementary Fig. 2**), including (1) Family 1: vmFPC-only or dmPFC-only inputs to the amygdala (models 1-6), (2) Family 2: both vmPFC and dmPFC inputs to the amygdala but with only one region connecting to the other two (models 7-18), (3) Family 3: both vmPFC and dmPFC inputs to the amygdala with all three regions connecting with each other unidirectionally (models 19-36), and (4) Family 4: both vmPFC and dmPFC inputs to the amygdala with at least two regions connecting with each other bidirectionally (models 37-44).

The Random-effects Bayesian Model Selection (RFX-BMS) procedure was used to determine the best model at the group level (i.e., we compared the model evidence for all 44 pre-defined models). RFX-BMS reports the posterior probability (i.e., how likely a specific model generates the data of a randomly chosen participant) and the exceedance probability (i.e., how likely a given model is more frequent than any other models). The RFX-BMS is itself a statistical inference (statement of relative probabilities) but not an index of the goodness of model fit for a dataset. In the RFX-BMS, models are treated as random effects that can differ between participants and have a fixed (unknown) distribution in that population. Given that the RFX-BMS does not assume that the optimal model is the most likely one for each individual participant and is therefore less susceptible to outliers in the experimental data than fixed-effects (FFX) methods [52, 53]. This procedure also implies that model selection is relativistic, i.e., it compares models against each other. Furthermore, the expected posterior and exceedance probabilities of an individual model decrease as the number of models increase. Hence, we only examined a set of highly plausible models based on our hypotheses and interpreted the model with the highest exceedance probability (i.e., the best model).

### Single-neuron electrophysiology and differential latency analysis

Detailed methods of single-neuron electrophysiology have been described in our previous study [5]. Briefly, we recorded bilaterally from implanted depth electrodes in the amygdala and dmPFC (dACC and pre-supplementary motor area [pre-SMA]). Only single units with an average firing rate of at least 0.2 Hz (entire task) were considered. Trials were aligned to stimulus onset. Average firing rates (PSTH) were computed by counting spikes across all trials in consecutive 250 ms bins.

For differential latency analysis, we binned spike trains into 1-ms bins and computed the cumulative sum. We then averaged the cumulative sums for each ambiguity level. We here only analyzed amygdala neurons that preferred unambiguous faces and dmPFC neurons that preferred ambiguous faces because amygdala neurons preferring ambiguous faces and dmPFC neurons preferring unambiguous faces had different temporal dynamics. We then compared, at every point of time, whether the cumulative sums of a group of neurons were different (P < 0.01, one-tailed pairwise *t*-test; FDR corrected). The first point of time of the significant cluster (cluster size > 10 time points) was used as the estimate of the differential latency. Note that this method is not sensitive to differences in baseline firing rate between neurons because the latency estimate is pairwise for each neuron individually. To assess statistical significance, we estimated the null distribution by first randomly shuffling the labels for groups and then repeated the above latency analysis. We used 1000 runs for the permutation analysis. We compared the observed latency difference between groups with this null distribution of latency difference to obtain p-values.

### Electroencephalogram (EEG)

Detailed methods of EEG data recording and preprocessing, event-related potential (ERP) analysis, time frequency analysis, and source localization have been described in our previous reports [4, 32]. Below, we describe the methods for functional connectivity analyses, which have not been reported in our previous studies.

Our previous work [4] revealed the strongest ERP response to emotion ambiguity at the parietal-central (Pz) electrode (i.e., the late positive potential [LPP], starting from 400 ms after stimulus onset and lasting for 300 ms), and this response can be source localized to the dmPFC and vmPFC. Here, using the Pz as the source channel, we set out to identify other channels that showed coordination with the Pz and further test how such coordination was modulated by ambiguity levels.

### EEG: cross-channel coherence

An event-related EEG cross-channel coherence analysis was performed to identify the cortical coordination responding to emotion ambiguity. EEG coherence is defined as the normalized cross-power spectrum of two signals recorded simultaneously from different electrodes: 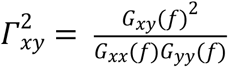, where *G_xy_*(*f*) is the cross-power spectral density and *G_xx_*(*f*) and *G_yy_*(*f*) are the respective auto-power spectral densities. EEG coherence is a measure of the consistency of relative amplitude between a pair of signals at a given frequency and can be interpreted as an index of their functional communication [54, 55]. EEG coherence was computed for all inter- and intra-hemispheric pairwise combinations of electrodes with the source electrode Pz, and it was computed for a combined frequency ranges from 4 to 23 Hz that mainly covered the theta, alpha, and beta frequency bands, given the duration of the LPP signal (300 ms). Notably, given that EEG coherence does not convey the direction of information flow, here, we only used the Pz as the source channel to search for its coherence with other channels instead of repeating the same analysis using other channels as the sources. Moreover, the magnitude-squared coherence is a function of frequency across channels whose values range from 0 to 1. A greater coherence value between a pair of electrodes indicates a greater synchronization between the electrodes.

### EEG: cross-frequency coupling

In addition to the functional connectivity at the channel level, we also examined the functional connectivity using cross-frequency coupling (CFC). The CFC denotes the interplay between two different frequencies, and phase/amplitude-amplitude CFC provides an effective means to integrate activity across different spatial and temporal scales [56, 57]. Phase/amplitude-amplitude CFC describes the statistical dependence between the phase/amplitude of a low-frequency brain rhythm and the amplitude of a high-frequency brain signal. Here, we focused on the lower-frequency delta band given its role in encoding emotion ambiguity [4] and its potential to couple with higher frequency bands (i.e., theta, alpha, and beta) based on a 2-second epoch (500 ms before stimulus onset to 1500 ms after stimulus onset). Specifically, we first filtered the data into high- and low-frequency bands. We then extracted the amplitude from the filtered signals. Lastly, we constructed GLMs (one for delta-band signal and one for theta/alpha/beta-band signal) to identify whether the amplitudes of the theta, alpha or beta signals varied as a function of the amplitude of the delta signal. Together, the amplitude-amplitude CFC could serve as the modulation index (or predictive value) of the delta amplitude on the theta, alpha, and beta amplitude.

### EEG: source connectivity

We computed the effective connectivity at the source level. Specifically, we used the group averaged LPP signals as inputs to identify the directed functional (“causal”) interactions among time-series data generated from the cortical sources. We first reconstructed the cortical source signals by implementing a cortical current density inverse imaging analysis using a realistic and spherical human head model. We then defined the ROI sources based on the reconstructed regions showing the strongest activation, which are generally consistent with the activated brain regions identified by fMRI [4]. Next, we applied a directed transfer function (DTF), a frequency-domain estimator of causal interaction based on the multivariate autoregressive (MVAR) modeling, to reveal the direction of the information flow among the cortical ROIs. We used a permutation test with 1000 runs to determine the statistical significance across the time course (300-600 ms after stimulus onset; in the frequency domain from 1 to 30 Hz). We implemented this analysis using the MATLAB toolbox eConnectome [58].

### Predicting functional connectivity in neuropsychiatric patients

We built a linear model to predict functional connectivity from behavioral response based on high-order PPI analysis:

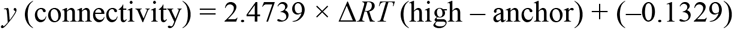

where *y* is the strength of amygdala-dmPFC connectivity, 2.4737 is the regression coefficient reflecting the correlation between RT difference (i.e., ambiguity sensitivity) and amygdala-dmPFC connectivity, and –0.1329 is the regression intercept. The regression parameters, 2.4739 and –0.1329, were fitted from the fMRI data used in high-order PPI analysis. A smaller RT difference suggests that participants were less sensitive to differences in emotion ambiguity in stimulus, which can lead to a smaller *y* that suggests a weaker amygdala-dmPFC connectivity; and vice versa for a greater RT difference and stronger amygdala-dmPFC connectivity. We then predicted *y* for each group of participants (neuropsychiatric or control) based on their RT difference.

### Data availability

All data is publicly available on OSF (https://osf.io/26rhz/).

## Results

### Behavior and functional localization

In the main experiment with fear-happy morphed faces, we asked participants to judge emotional faces as fearful or happy (**Fig. 1a**). Faces were either unambiguously happy, unambiguously fearful, or graded ambiguous morphs between the two emotions (**Fig. 1b**). Since emotion ambiguity was distributed symmetrically between the two emotions, we grouped the seven emotion levels into three ambiguity levels (**Fig. 1b**): anchor/unambiguous, intermediate (30%/70% morph), and high (40%-60% morph). Detailed behavioral quantification has been described in our previous reports [4, 5]. Briefly, for each participant, we quantified behavior as the proportion of trials identified as fearful as a function of morph level (**Fig. 1c, d**). We found a monotonically increasing relationship between the likelihood of identifying a face as fearful and the fearfulness in the morphed face. Both EEG and fMRI participants had similar psychometric curves (point-by-point comparison using *t*-test, corrected for false discovery rate (FDR) for Q < 0.05 [59]; all Ps > 0.05). Moreover, participants responded slower (relative to stimulus onset) for the faces with high ambiguous expressions compared to faces with unambiguous faces (**Fig. 1e, f**; one-way repeated measures ANOVA: *F*(6, 246) = 26.60, P < 10^-10^, η_p_^2^ = 1; *F*(2, 82) = 48.54, P = 1.23×10^-14^, η_p_^2^ = 1).

**Fig. 1.**
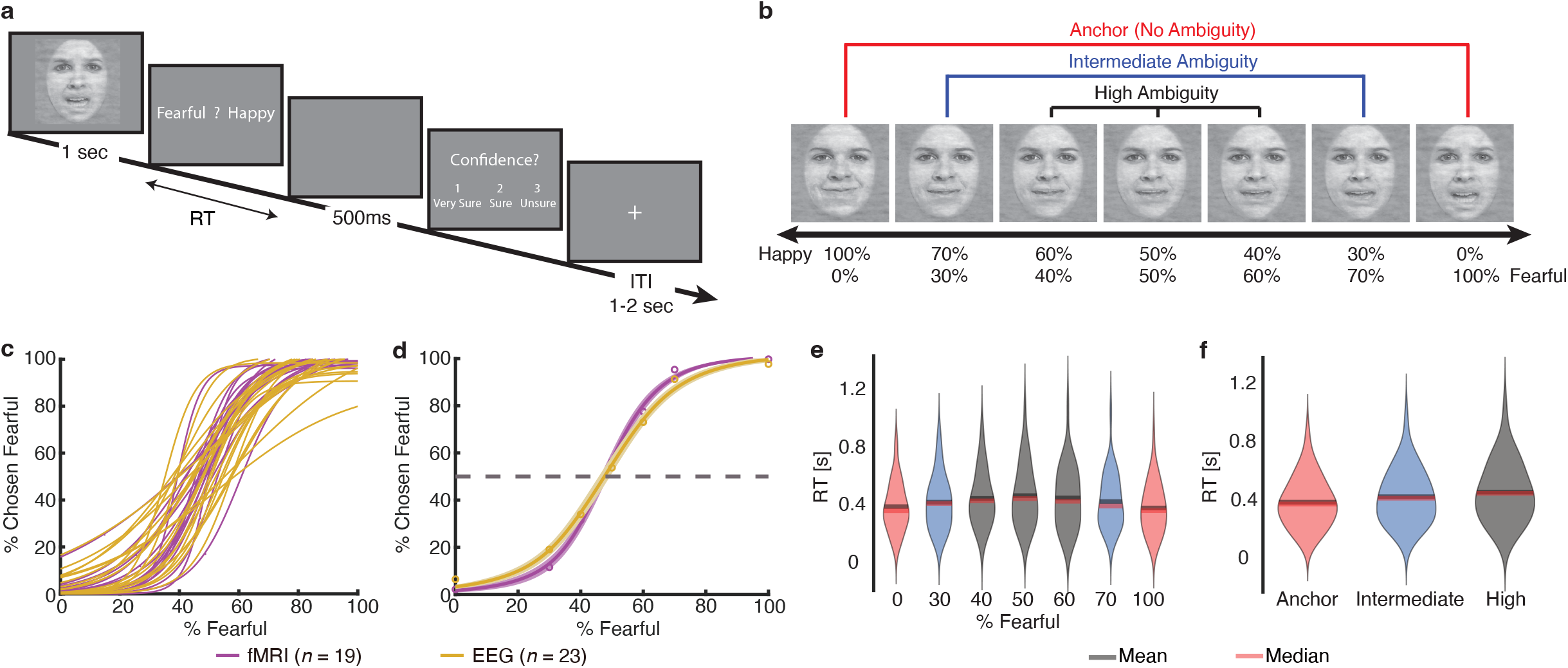
Behavior. **(a)** Task. A face was presented for 1 second followed by a question asking participants to identify the facial emotion (fearful or happy). The faces were chosen from the STOIC database, a database of face images expressing highly recognizable emotions, in which their facial expressions were morphed by a computer algorithm. For behavioral participants, after a blank screen of 500 ms, they were then asked to indicate their confidence in their decision (‘1 ‘for ‘very sure’, ‘2 ‘for ‘sure ‘or ‘3 ‘for ‘unsure’). Faces are not shown to scale. **(b)** Sample stimuli of one female identity ranging from 100% happy/0% fearful to 0% happy/100% fearful. Three ambiguity levels (unambiguous, intermediate, and high) are grouped as shown above the stimuli. **(c-f)** Behavioral results. **(c)** Psychometric curves from individual participants showing the proportion of trials judged as fearful as a function of morph levels (ranging from 0% fearful [100% happy; on the left] to 100% fearful [0% happy; on the right]). **(d)** Group average of psychometric curves. Shaded area denotes ±SEM across participants. **(e)** Reaction time (RT; relative to stimulus onset) for the fear/happy decision as a function of fearful level. **(f)** RT as a function of ambiguity level. Violin plots present the distribution of RT for combined fMRI and EEG participants (*n* = 42).

### PPI: PFC-amygdala connectivity is involved in encoding emotion ambiguity

Functional localization has revealed brain activation to emotion ambiguity: we found a significant increase of BOLD signal in the bilateral dmPFC and inferior frontal gyrus (IFG)/anterior insula with increasing level of emotion ambiguity (**Fig. 2a**) and a significant increase of BOLD signal in the right amygdala, left vmPFC, posterior cingulate cortex (PCC), dorsolateral prefrontal cortex (dlPFC), inferior parietal lobule (IPL), and right postcentral gyrus with decreasing level of emotion ambiguity (**Fig. 2b**). Based on these functional localization results, we next investigated the relationships between these brain regions involved in encoding emotion ambiguity.

**Fig. 2.**
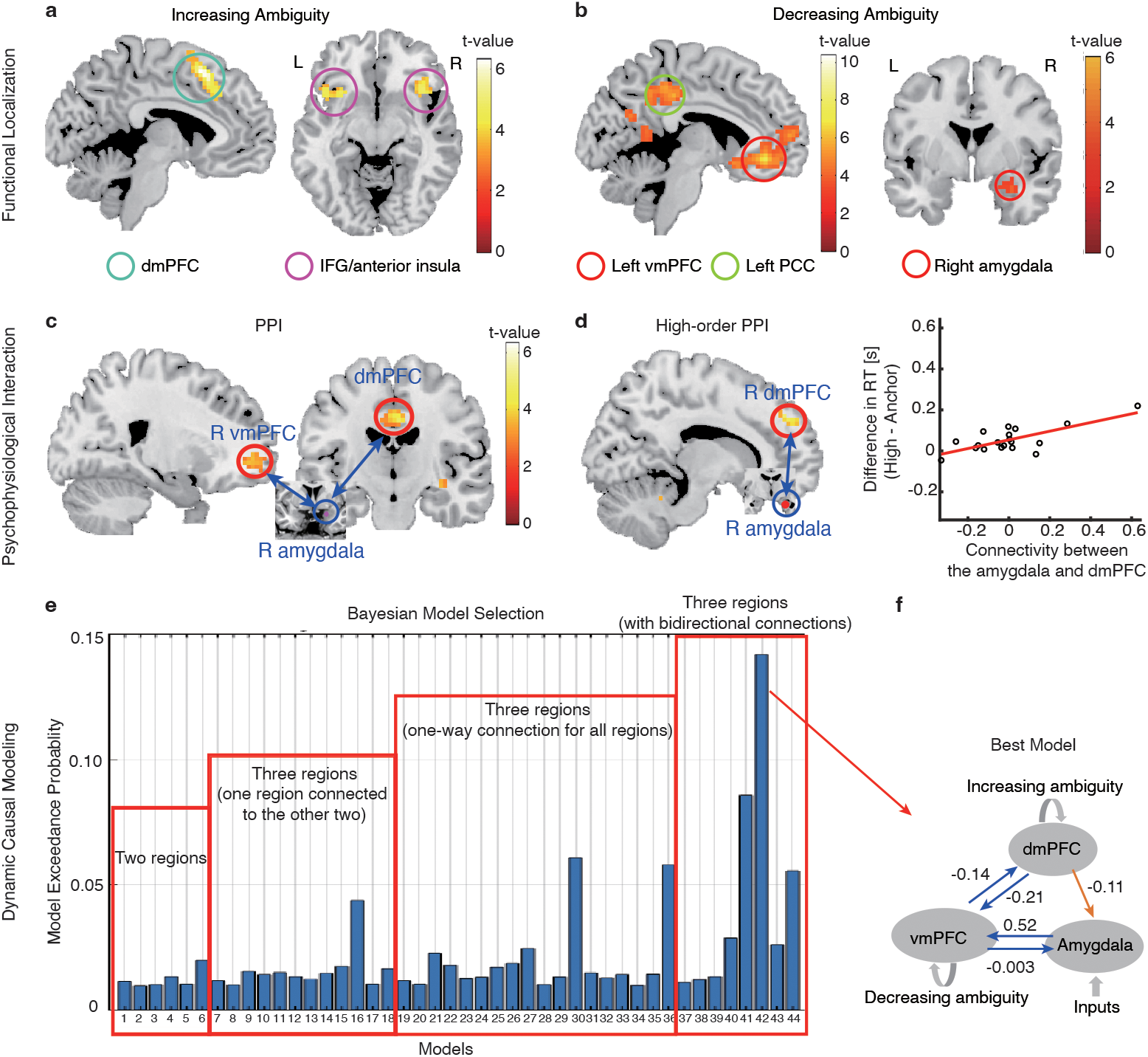
Psychophysiological interaction (PPI) and dynamic causal modeling (DCM). **(a, b)** Functional localization. **(a)** Increasing ambiguity was correlated with increasing BOLD activity in the bilateral dorsomedial prefrontal cortex (dmPFC) and inferior frontal gyrus (IFG)/anterior insula. The generated statistical parametric map was superimposed on anatomical sections of the standardized MNI T1-weighted brain template. L: left. R: right. **(b)** Decreasing ambiguity was correlated with increasing BOLD activity in the right amygdala, left ventromedial prefrontal cortex (vmPFC), and posterior cingulate cortex (PCC). **(c)** PPI analysis revealed functional connectivity between the amygdala and vmPFC as well as between the amygdala and dmPFC. **(d)** High-order PPI revealed functional connectivity between the amygdala and dmPFC. In the right plot, each dot represents a participant (*n* = 19), and the red line represents the linear fit. **(e)** Exceedance probability for each individual DCM model. Individual models are grouped into families (shown in red rectangles). **(f)** The best model shows a bidirectional connection between the vmPFC and amygdala, a bidirectional connection between the dmPFC and vmPFC, and a unidirectional connection from the dmPFC to the amygdala. Numbers show the mean of the maximum a posteriori (MAP) estimates of the optimal model.

We employed the PPI analysis to identify target regions that were functionally connected to the source (i.e., the right amygdala; *x* = 30, *y* = 0, *z* = –21) and whose connectivity was further modulated by ambiguity levels (see **Methods**). We found that the right vmPFC (**Fig. 2c**; peak: *x* = 21, *y* = 48, *z* = –3; 24 voxels, SVC, FWE P = 0.05) and bilateral dmPFC (**Fig. 2c**; peak: *x* = 0, *y* = –15, *z* = 39; 14 voxels, SVC, FWE P = 0.014) were positively correlated with the activity in the amygdala, suggesting an amygdala-PFC functional network that encoded emotion ambiguity. Notably, this amygdala-PFC network (especially the left amygdala) was also engaged in encoding emotion intensity (**Supplementary Fig. 1**).

We next employed high-order PPI models (see **Methods**) to identify brain regions whose functional connectivity with the right amygdala correlated with behavioral ambiguity sensitivity (i.e., difference in reaction times between high ambiguity and anchor conditions). We found that the functional connectivity between the right amygdala and right dmPFC (**Fig. 2d**; peak: *x* = 12, *y* = 36, *z* = 48; 26 voxels, P < 0.001 uncorrected) was positively correlated with the increasing ambiguity sensitivity (**Fig. 2d**), suggesting that inter-participant variability in ambiguity sensitivity modulated the amygdala-PFC functional connectivity.

### DCM: directional effective connectivity involved in encoding emotion ambiguity

Our PPI results have primarily identified two brain regions, the vmPFC and dmPFC, that are functionally connected with the amygdala. However, no directional information was shown in the PPI results. To further pinpoint the directional pathway among these three brain regions, we performed a DCM analysis. The index derived from the RFX-BMS, model exceedance probability, was used to evaluate the DCM models (see **Methods**); and the model with the highest exceedance probability was considered as the best model [60–63]. Overall, the models from Family 4 showed a higher exceedance probability than the models from other families (**Fig. 2e**). In particular, the model with a bidirectional connection between the vmPFC and amygdala, a bidirectional connection between the dmPFC and vmPFC, and a unidirectional connection from the dmPFC to the amygdala, outperformed all other models with an exceedance probability of approximately 15% (**Fig. 2f**; see other models in **Supplementary Fig. 2**). The bidirectional connectivity between the vmPFC and amygdala may reflect processes involving both emotion appraisal (i.e., amygdala➔vmPFC➔dmPFC) and emotion regulation (i.e., dmPFC➔vmPFC➔amygdala). Using brain areas all from the right hemisphere (**Supplementary Fig. 3**), we not only replicated the top-down connectivity from the dmPFC to the amygdala and the bottom-up connectivity through the vmPFC, but also revealed a direct bottom-up connectivity from the amygdala to the ipsilateral dmPFC. Together, our results suggest both bottom-up affective processes for ambiguity representation/perception and topdown cognitive processes for ambiguity resolution/decision.

### Single-neuron differential latency analysis

To further elucidate the relationship between the amygdala and dmPFC, we recorded single-neuron activity from the amygdala and dmPFC (dACC and pre-SMA) from 16 neurosurgical patients. We recorded from 321 neurons in the amygdala (21 sessions) and 236 neurons in the dmPFC (15 sessions; overall firing rate greater than 0.2 Hz). It is worth noting that the recording locations were in the vicinity of the areas showing BOLD activation.

We investigated whether the responses of amygdala and dmPFC neurons were modulated by the level of emotion ambiguity. We used a linear regression to identify neurons whose firing rate correlated trial-by-trial with three levels of emotion ambiguity. We found 36 amygdala neurons (11.2%; binomial P = 2.58×10^-6^; see **Fig. 3a** for an example and **Fig. 3c, e** for group result) and 29 dmPFC neurons (12.3%; binomial P = 3.09×10^-6^; see **Fig. 3b** for an example and **Fig. 3d, f** for group result) that showed a significant trial-by-trial correlation. Notably, consistent with the fMRI results (**Fig. 2a, b**), the majority of amygdala neurons (33/36; χ^2^-test: P = 1.54×10^-12^) had the maximal firing rate for unambiguous faces whereas the majority of dmPFC neurons (18/29; χ^2^-test: P = 0.066) had the maximal firing rate for the most ambiguous faces.

**Fig. 3.**
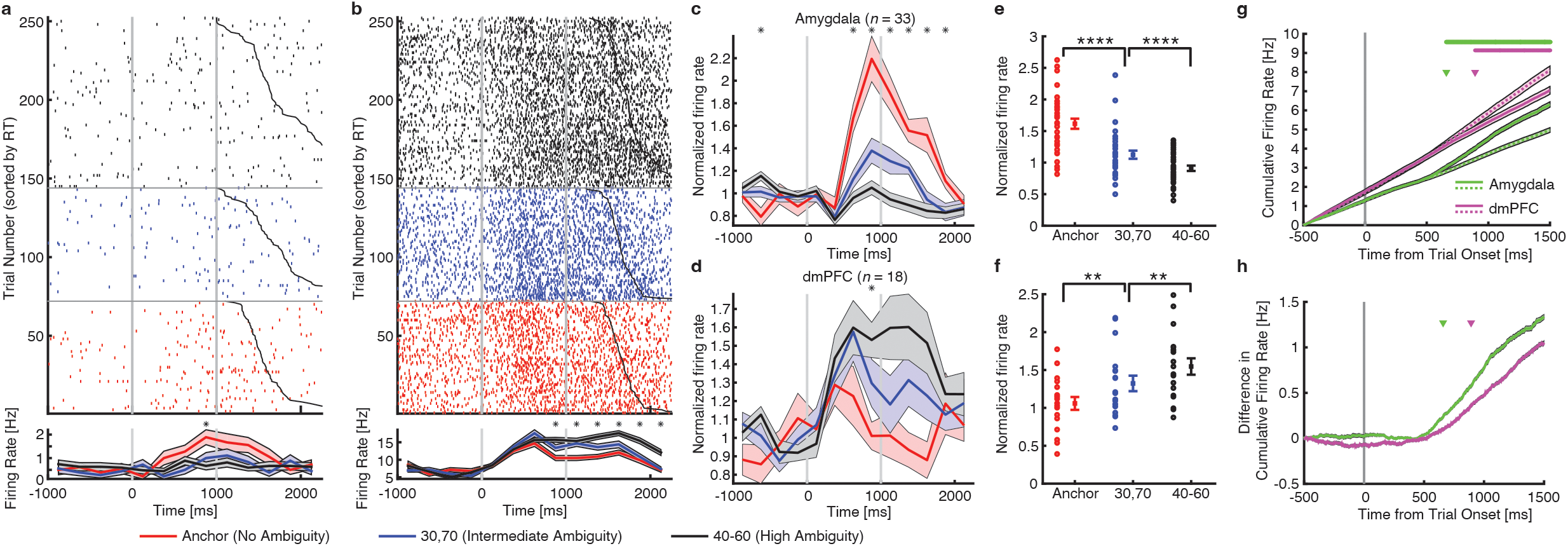
Single-neuron differential latency. **(a)** An example amygdala neuron that fire most to unambiguous faces and least to the most ambiguous faces (linear regression: P < 0.05). **(b)** An example dorsomedial prefrontal cortex (dmPFC) neuron that fire most to the most ambiguous faces and least to unambiguous faces (linear regression: P < 0.05). Raster (top) and PSTH (bottom) are color coded according to ambiguity levels as indicated. Trials are aligned to face stimulus onset (left gray bar, fixed 1s duration) and sorted by reaction time (black line). PSTH bin size is 250 ms. Shaded area and error bars denote ±SEM across trials. Asterisk indicates a significant difference between the conditions in that bin (P < 0.05, one-way ANOVA, Bonferroni-corrected). **(c, d)** Average normalized firing rate of ambiguity-coding neurons. Asterisk indicates a significant difference between the conditions in that bin (P < 0.05, oneway ANOVA, Bonferroni-corrected). **(e, f)** Mean normalized firing rate at ambiguity level. Normalized firing rate for each unit (left) and mean±SEM across units (right) are shown at each ambiguity level. Mean firing rate was calculated in a time window 250 to 1750 ms after stimulus onset (the same time window as neuron selections). Asterisks indicate a significant difference between conditions using paired two-tailed *t*-test. **: P < 0.01 and ****: P < 0.0001. **(c, e)** Neurons in the amygdala that increased their firing rate for the least ambiguous faces (*n* = 33). **(d, f)** Neurons in the dmPFC that increased their firing rate for the most ambiguous faces (*n* = 18). **(g)** Cumulative firing rate for neurons from the amygdala (green lines; *n* = 36 neurons) and dmPFC (magenta lines; *n* = 29 neurons). Shaded area denotes ±SEM across neurons. Solid lines: unambiguous faces. Dotted lines: the most ambiguous faces. Top bars show clusters of time points with a significant difference (one-tailed pairwise *t*-test; P < 0.01; FDR-corrected; cluster size > 10 time points). Arrows indicate the first time point of the significant cluster. Green: amygdala neurons. Magenta: dmPFC neurons. **(h)** Difference in cumulative firing rate (same data as shown in **(g)**). Shaded area denotes ±SEM across neurons. Arrows indicate the first time point of the significant cluster. Green: amygdala neurons. Magenta: dmPFC neurons.

We next compared the onset latency, relative to stimulus onset, of the ambiguity-coding neurons between the amygdala and dmPFC. We found that amygdala ambiguity-coding neurons (*n* = 36) responded significantly earlier than dmPFC ambiguity-coding neurons (*n* = 29; **Fig. 3g, h**; amygdala: 658 ms relative to stimulus onset; dmPFC: 893 ms; permutation test: P = 0.045; such difference in onset latency can be appreciated from the single-neuron examples shown in **Fig. 3a, b**). This result was similar for dACC (permutation test: P < 0.001) and pre-SMA neurons (permutation test: P < 0.001). Together, this latency difference reflects the bottom-up process for ambiguity processing and is consistent with the pathway revealed by DCM analysis (amygdala➔vmPFC➔dmPFC).

### EEG cross-channel coherence and cross-frequency coupling

Our prior EEG results have complemented the fMRI findings by providing a higher temporal resolution and the single-neuron findings by providing a broader spatial coverage [4]: (1) the late positive potential (LPP) encodes levels of emotion ambiguity, (2) neural oscillation in the delta frequency bands correlates with emotion ambiguity, and (3) regional sources of the LPP are highly consistent with the brain regions showing a significant increase of BOLD signal. In addition, we have conducted a series of control experiments to elucidate the role of the LPP in encoding perceptual ambiguity [4, 32]. Here, we further examined the functional neural network based on scalp electrophysiology.

First, we performed a cross-channel coherence analysis to identify the channels that were functionally connected with the channel Pz (the channel showing the LPP) during ambiguity processing. We found that the central region (shown by the activity from the Cz electrode) was functionally connected with the parietal-central region (shown by the activity from the Pz electrode) during ambiguity processing (**Fig. 4a**). Such connectivity varied as a function of emotion ambiguity, with unambiguous stimuli eliciting the strongest coherence (**Fig. 4a**; oneway repeated measures ANOVA: *F*(2, 42) = 3.58, P = 0.036, η_p_^2^ = 0.63). Therefore, parietal-central coherence was involved in encoding emotion ambiguity. Notably, we replicated this finding in the task with a speeded response: the central (Cz) and frontal-central (FCz) regions were connected with the parietal-central region (**Fig. 4b**; *F*(2, 30) = 3.95, P = 0.029, η_p_^2^ = 0.66), which further confirmed that parietal-central EEG coherence was involved in encoding emotion ambiguity.

**Fig. 4.**
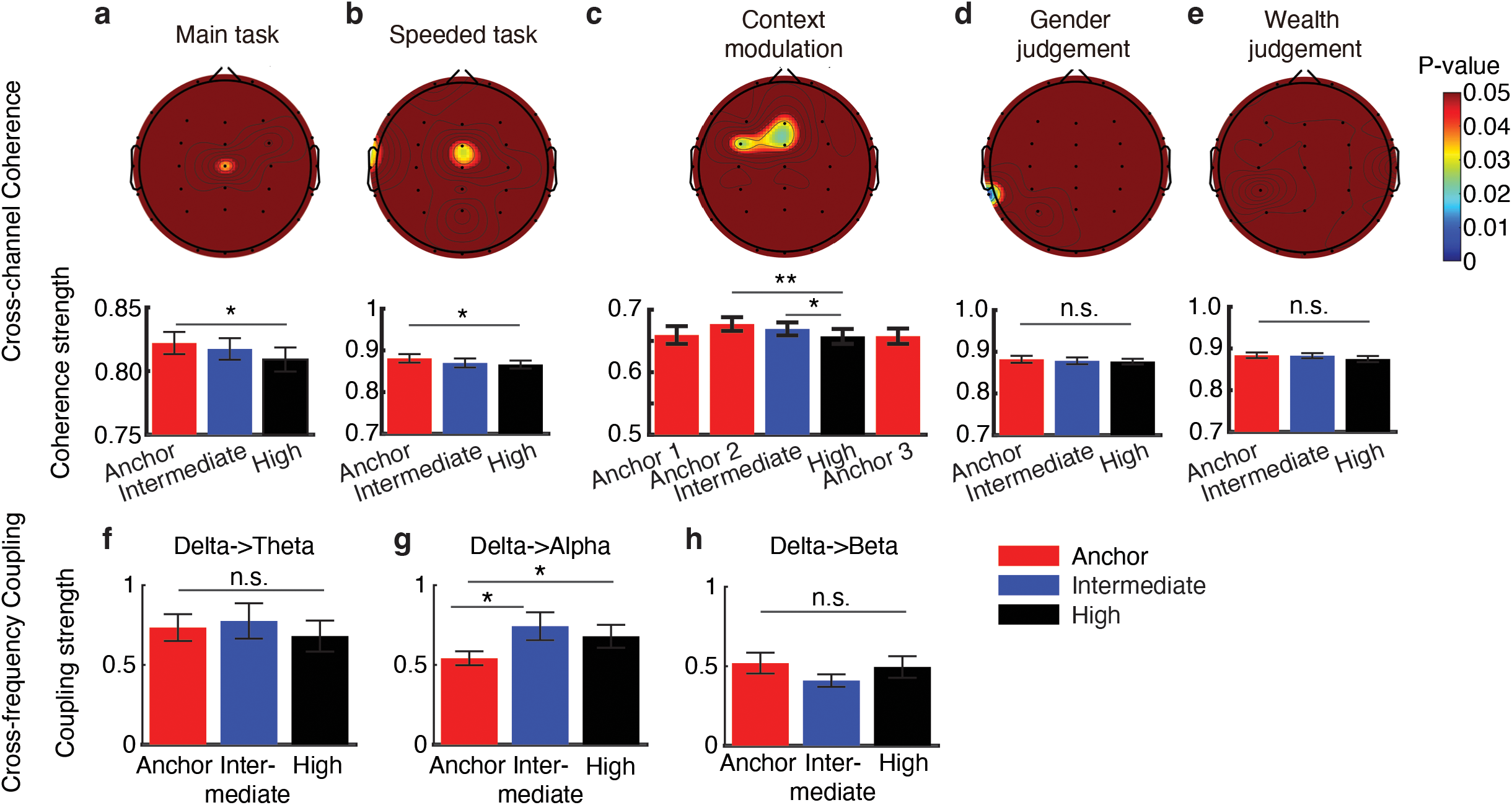
EEG cross-channel coherence and cross-frequency coupling. **(a-e)** Cross-channel coherence. **(a)** Main task. **(b)** Task with a speeded response. **(c)** Task with a context modulation. Anchor 1: anchor faces from the first block. Anchor 2: anchor faces from the second block. Anchor 3: anchor faces from the third block. **(d)** Gender judgment task. **(e)** Wealth judgment task. Color coding in the topography indicates p-values (for P < 0.05 only). Error bars denote one SEM across participants. Asterisks indicate a significant difference across conditions using one-way repeated measures ANOVA. *: P < 0.05 and **: P < 0.01. n.s.: not significant (P > 0.05). **(f-h)** Cross-frequency coupling. **(f)** Delta-theta coupling. **(g)** Delta-alpha coupling. **(h)** Delta-beta coupling. Error bars denote one SEM across participants. Asterisks indicate a significant difference between/across conditions using two-tailed paired *t*-test or one-way repeated measures ANOVA. *: P < 0.05. n.s.: not significant (P > 0.05).

We next conducted two additional experiments to investigate the modulation of EEG cross-channel coherence. In a task with different contexts (i.e., whether ambiguous faces were present with anchor faces; see **Methods**), we first found in the second block that the central (Cz) and frontal-central (FCz) regions were connected with the parietal-central region (**Fig. 4c**; *F*(2, 56) = 5.99, P = 0.004, η_p_^2^ = 0.86), again replicating the finding in our main experiment (**Fig. 4a**). By comparing the coherence for anchor faces in the first, second, and third blocks, we found a reduced coherence when no ambiguous stimuli were present (**Fig. 4c**; *F*(2, 56) = 4.48, P = 0.043, η_p_^2^ = 0.53), indicating that the parietal-central coherence was modulated by the context of ambiguous stimuli. However, the parietal-central coherence was abolished when the judgment decision was certain (judging the gender of the face model; **Fig. 4d**; P > 0.1) or when the judgment decision was not congruent with stimulus ambiguity (judging the wealth [poor versus rich] of the face model; **Fig. 4e**; P > 0.1). Therefore, the parietal-central coherence was specific to decisions made on a dimension that was ambiguous.

Lastly, we investigated the functional connectivity in the spectral domain. We found that the alpha power was modulated by the delta phase, and their connectivity varied as a function of emotion ambiguity, with the most ambiguous stimuli eliciting the strongest delta-alpha cross-frequency coupling (**Fig. 4g**; *F*(2, 42) = 4.03, P = 0.025, η_p_^2^ = 0.68). However, no significant difference was found for the delta-theta (**Fig. 4f**; *F*(2, 42) = 0.41, P = 0.66, η_p_^2^ = 0.11) or the delta-beta (**Fig. 4h**; *F*(2, 42) = 1.72, P = 0.19, η_p_^2^ = 0.34) cross-frequency coupling.

Together, our results revealed parietal-frontal-central cross-channel coherence as well as delta-alpha cross-frequency coupling in processing emotion ambiguity.

### EEG source connectivity: the dmPFC top-down regulates the activities in other brain regions

We next investigated the effective connectivity in a source domain using the directed transfer function, which could localize the origins of EEG signals and their directional connections. Six brain regions were covered in the effective connectivity analysis, including the right dmPFC, bilateral superior frontal gyrus (SFG), right vmPFC, bilateral dlPFC, bilateral IPL, and left occipital cortex (**Fig. 5a**). We found that the right dmPFC had a directed information flow (i.e., connectivity) to the left SFG, right dlPFC, right vmPFC, and right IPL (**Fig. 5a, b**), suggesting a top-down modulation of brain activity during processing of emotion ambiguity. Notably, the directed source connectivity from the right dmPFC to the right vmPFC was consistent with our DCM results where we demonstrated a bidirectional connectivity between these brain regions.

**Fig. 5.**
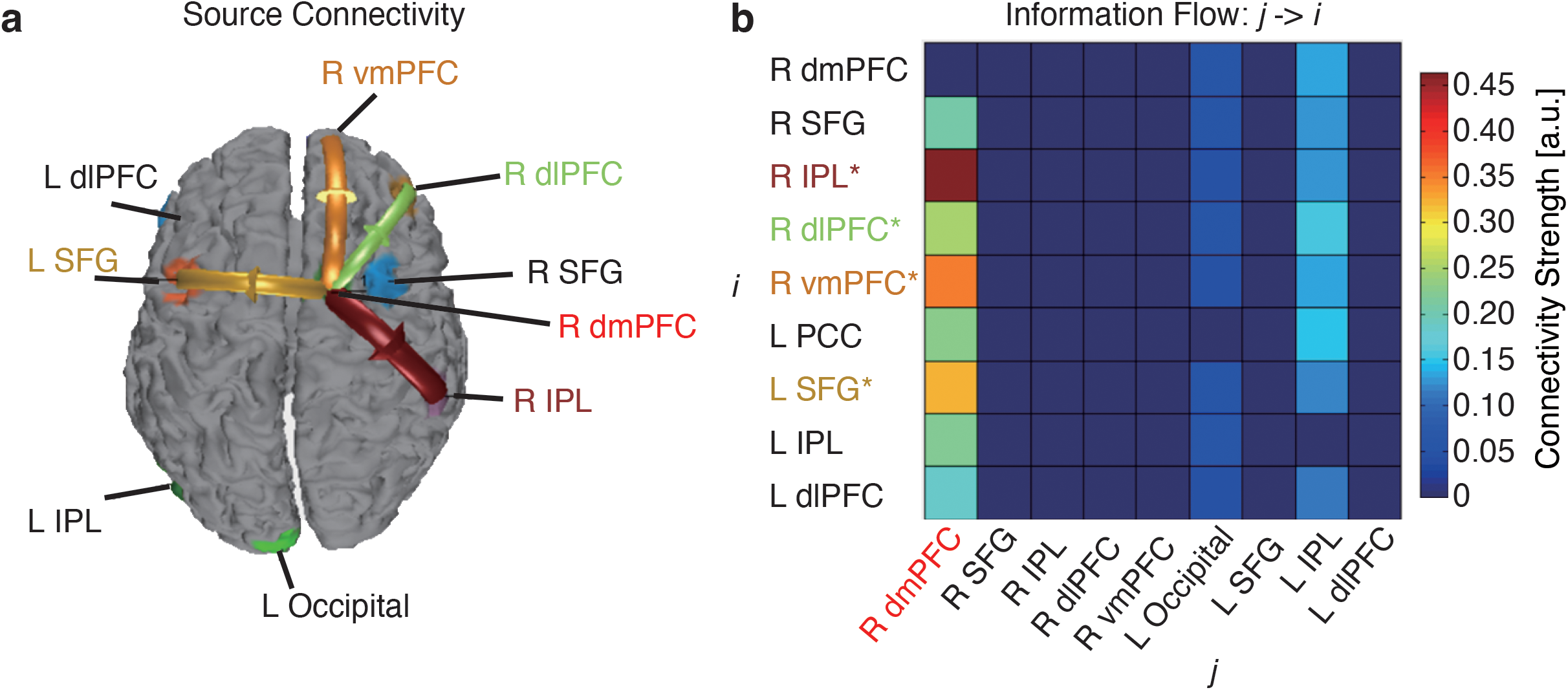
EEG source connectivity. **(a)** Identified source connectivity. **(b)** Connectivity matrix. Color coding in the connectivity matrix indicates the strength of information flow from source *j* to source *i*. Asterisks indicate brain regions with a significant directional information flow from the dmPFC (P < 0.05; i.e., brain regions showing a significant connectivity with the dmPFC). L: left. R: right. dmPFC: dorsomedial prefrontal cortex. SFG: superior frontal gyrus. IPL: inferior parietal lobule. dlPFC: dorsolateral prefrontal cortex. vmPFC: ventromedial prefrontal cortex.

### Behavioral deficits in neuropsychiatric patients

Above, we have systematically investigated the functional connectivity during processing of emotion ambiguity using multimodal approaches. Previous research has shown intrinsic or task-induced dysfunction of the amygdala-PFC network in neuropsychiatry patients with anxiety [39, 40], depression [41–43], ASD [36, 37], ADHD [38], and schizophrenia (SCZ) [45]. However, it remains unclear whether these patients show behavioral deficits in processing perceptual ambiguity of facial expressions. To answer this question, here we performed the same main task in a group of in-lab participants (**Fig. 6a-f**) and a group of online participants (**Fig. 6g-l**).

**Fig. 6.**
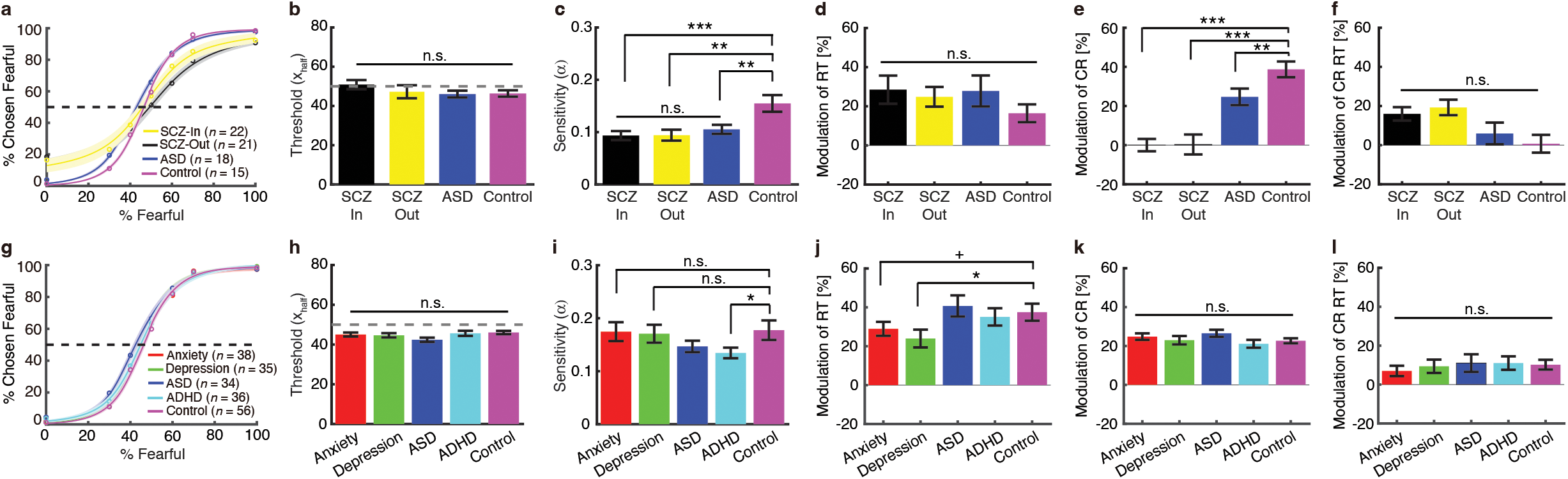
Behavioral results from neuropsychiatric patients. **(a-f)** In-lab patients with confirmed diagnosis of neuropsychiatric disorders. **(g-l)** Online patients with self-reported neuropsychiatric disorders. **(a, g)** Group average of psychometric curves. Legend conventions as in **Fig. 1**. **(b, h)** Emotion discrimination threshold (*xhalf*). **(c, i)** Sensitivity to emotion intensity (*α*). **(d, j)** Modulation of emotion judgment reaction time (RT). **(e, k)** Modulation of confidence rating (CR). **(f, l)** Modulation of confidence rating reaction time (CR RT). Modulation was defined as the difference between high ambiguity and anchor (high - anchor), normalized by the response to anchor. Asterisks indicate a significant difference between groups using two-tailed two-sample *t*-test. +: P < 0.1, *: P < 0.05, **: P < 0.01, and ***: P < 0.001. n.s.: not significant.

We first examined in-lab patients with confirmed diagnosis of neuropsychiatric disorders (including in-patient SCZ, out-patient SCZ, and ASD) and controls. Although we did not find a significant difference in emotion discrimination threshold in patients (**Fig. 6a, b**; *F*(3, 75) = 0.83, P = 0.47, η_p_^2^ = 0.09), we found reduced sensitivity to emotion intensity in each patient group (**Fig. 6a, c**; *F*(3, 75) = 6.40, P = 0.001, η_p_^2^ = 0.73; in-patient SCZ vs. control: *t*(35) = 3.66, P = 0.001, *d* = 0.61; out-patient SCZ vs. control: *t*(34) = 3.32, P = 0.002, *d* = 0.56; ASD vs. control: *t*(31) = 2.85, P = 0.008, *d* = 0.51). Reduced sensitivity in emotion intensity (i.e., flatter psychometric curves and smaller *α*) suggest that both SCZ and ASD patients were less specific in their emotion judgments, since they made similar judgments given different morph levels. Furthermore, although patients did not show a significant difference in the modulation (high - anchor) of emotion judgment RT (**Fig. 6d**; *F*(3, 75) = 0.55, P = 0.64, η_p_^2^ = 0.06) or confidence rating RT (**Fig. 6f**; *F*(3, 64) = 0.35, P = 0.64, η_p_^2^ = 0.04), all patient groups demonstrated a significantly reduced modulation of confidence rating (**Fig. 6e**; *F*(3, 64) = 5.84, P = 0.0014, η_p_^2^ = 0.73; in-patient SCZ vs. control: *t*(31) = 4.35, P = 1.37×10^-4^; out-patient SCZ vs. control: *t*(30) = 3.13, P = 0.0039, *d* = 0.57; ASD vs. control: *t*(20) = 2.40, P = 0.026, *d* = 0.53; see data for each condition in **Supplementary Fig. 4**). The RT for the fear/happy decision can be considered as an implicit measure of confidence. Therefore, patients with SCZ and ASD only demonstrated deficits in explicit (**Fig. 6e**) rather than implicit (**Fig. 6d**) confidence judgments.

We also explored a group of online patients with self-reported diagnosis of neuropsychiatric disorders (including anxiety, depression, ASD, and ADHD) and controls. We found no significant difference in emotion discrimination threshold in patients (**Fig. 6g, h**; *F*(4, 198) = 1.79, P = 0.13, η_p_^2^ = 0.13). Compared to the controls, we found reduced sensitivity to emotion intensity in patients with ADHD (**Fig. 6g, i**; *t*(90) = 2.05, P = 0.044, *d* = 0.21), a trend in patients with ASD (*t*(88) = 1.43, P = 0.15, *d* = 0.15; cf. **Fig. 6c**), but not in patients with anxiety (*t*(92) = 0.10, P = 0.91, *d* = 0.01) or depression (*t*(89) = 0.25, P = 0.80, *d* = 0.02). Moreover, we observed a significantly reduced modulation of emotion judgment RT in patients with depression (**Fig. 6j**; *F*(4, 198) = 3.55, P = 0.031, η_p_^2^ = 0.25; depression vs. control: *t*(89) = 2.33, P = 0.021, *d* = 0.25), a trend in patients with anxiety (*t*(92) = 1.86, P = 0.065, *d* = 0.19), but not in patients with ASD or ADHD (both Ps > 0.05; see data for each condition in **Supplementary Fig. 4**). On the other hand, for all patient groups, we did not observe a significant difference in the modulation of confidence rating (**Fig. 6k**; *F*(4, 198) = 1.29, P = 0.27, η_p_^2^ = 0.09) or confidence rating RT (**Fig. 6l**; *F*(4, 198) = 0.25, P = 0.91, η_p_^2^ = 0.01).

Together, we surveyed both in-lab and online patients with neuropsychiatric disorders who have dysfunctions in amygdala-PFC functional connectivity and revealed altered behavioral responses in several aspects of emotion judgment in these patients.

### Predicting functional connectivity in neuropsychiatric patients

We lastly explored whether behavioral deficits in neuropsychiatric patients could be reflected in the amygdala-dmPFC connectivity. To address this question, we built a predictive model based on high-order PPI (i.e., the linear relationship between behavioral response to ambiguity and amygdala-dmPFC connectivity; **Fig. 2d**). We derived model parameters from fMRI participants and then predicted functional connectivity for each group of participants based on this model (see **Methods**). For in-lab participants (**Fig. 7a**), compared to controls, we found *enhanced* amygdala-dmPFC connectivity in in-patient SCZ patients (two-tailed two-sample *t*-test: *t*(35) = 3.32, P = 0.0021, *d* = 0.56) and out-patient SCZ patients (*t*(34) = 3.50, P = 0.0013, *d* = 0.61), but not in ASD patients (*t*(31) = 0.89, P = 0.37, *d* = 0.16). For online participants (**Fig. 7b**), compared to controls, we found *reduced* amygdala-dmPFC connectivity in patients with depression (*t*(89) = –2.17, P = 0.033, *d* = 0.23), but not in patients with anxiety (*t*(92) = – 1.55, P = 0.12, *d* = –0.16), ASD (*t*(88) = 0.14, P = 0.89, *d* = 0.015), or ADHD (*t*(90) = –0.39, P = 0.70, *d* = –0.04). Together, our results suggest that behavioral deficits in emotion judgment can be translated into abnormal functional connectivity in neuropsychiatric patients.

**Fig. 7.**
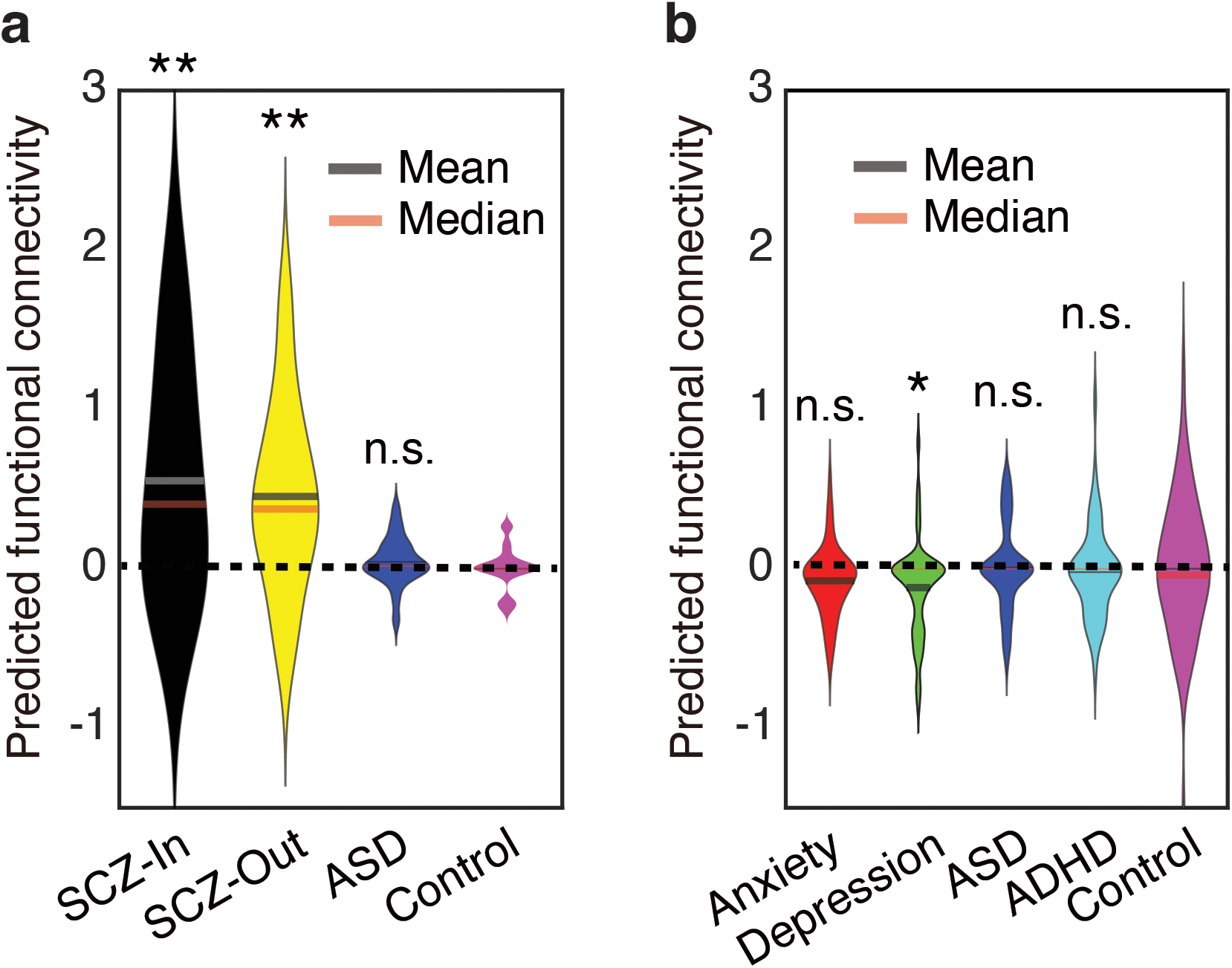
Predicted functional connectivity. **(a)** In-lab patients with confirmed diagnosis of neuropsychiatric disorders. **(b)** Online patients with self-reported neuropsychiatric disorders. Violin plots present the distribution of predicted functional connectivity (normalized by subtracting the mean of the controls). Asterisks indicate a significant difference compared to controls using two-tailed two-sample *t*-test. *: P < 0.05 and **: P < 0.01. n.s.: not significant.

## Discussion

Motivated by our prior studies showing that (1) both single neurons and BOLD-fMRI in the human amygdala parametrically encode levels of emotion ambiguity [5], (2) there is vast activation of the dmPFC and vmPFC for emotion ambiguity [4], and (3) the LPP originating from the dmPFC and vmPFC differentiates levels of emotion ambiguity and mediates behavioral judgments about ambiguous choices [4, 32], in the present study, we employed multimodal functional connectivity analyses to study the neural network underlying perceiving and resolving emotion ambiguity. PPI analysis showed amygdala-PFC connectivity in encoding emotion ambiguity, and DCM analysis revealed the directional effective connectivity between the amygdala, dmPFC, and vmPFC in this process. Furthermore, the responses of amygdala and dmPFC neurons were modulated by the level of emotion ambiguity, and amygdala neurons responded earlier than dmPFC neurons. We further found parietal-frontal coherence and delta-alpha cross-frequency coupling involved in encoding emotion ambiguity. In addition, EEG source connectivity revealed that the dmPFC top-down regulated the activities in other brain regions. Therefore, we have derived a network-level understanding of how the brain processes ambiguous facial expressions. We lastly showed altered behavioral responses in several groups of patients who have dysfunctions in amygdala-PFC functional connectivity.

### The amygdala-PFC network for processing emotion ambiguity

Emotional stimuli activate a broad network of brain regions, including the amygdala, dmPFC, and vmPFC. In humans, there has been increasing evidence showing that functional connectivity between the PFC and amygdala is critical for processing facial emotions. Our present results have pointed to an emotion processing circuit that underlie representing and resolving emotion ambiguity, consistent with convergent data from lesion, electrophysiology, and imaging studies showing that the PFC interacts with the amygdala to regulate cognitive and emotional processing [64–66]. Specifically, the amygdala may encode and represent the content of the emotion [5, 7] and provide input to the PFC. The dmPFC has been associated with cognitive processing and the vmPFC has been associated with affective processing [28]. The affective division of the PFC (i.e., vmPFC) modulates autonomic activity and internal emotional responses, while the cognitive division (i.e., dmPFC) is engaged in action selection associated with skeletomotor activity and motor response [65, 66]. The response in the dmPFC may thus reflect a control signal that resolves emotion ambiguity, consistent with our prior results that the LPP originating from the dmPFC is specifically associated with decisions (rather than stimulus) about ambiguity [4].

Furthermore, the amygdala may also reflect competition between passive and active responses to aversive stimuli [67], punishment predictions or prediction errors [68], or a more general source of information about errors [69], which recruits both cognitive (dmPFC) and emotional (amygdala) monitoring systems [69]. Such amygdala signals can be conveyed directly to the dmPFC or indirectly through connections from the amygdala to the striatum, insula, or vmPFC [22], consistent with the pathways identified in the present study. Furthermore, consistent with our DCM results of negative modulation of amygdala activity, previous analysis has revealed that both dmPFC and vmPFC show an inverse interaction with the direct thalamus-amygdala pathway [70].

### Atypical functional connectivity in neuropsychiatric disorders

A breakdown in the amygdala-PFC functional/effective connectivity may give rise to a variety of emotion-related deficits seen in a wide range of neuropsychiatric disorders. For example, enhanced amygdala-vmPFC bottom-up effects have been observed in major depressive disorders when processing emotional faces [42]. Such alterations can be even observed in infants exposed to prenatal maternal depression [71]. Moreover, genetic variations in the monoamine oxidase A (MAOA) associated with reduced amygdala-PFC coupling can predict the course and severity of major depression [72]. In addition, abnormal amygdala-PFC effective connectivity to happy faces differentiates bipolar from major depression [41], and antidepressant treatment efficacy can be measured by a significantly increased coupling between the amygdala and right PFC [73]. Consistent with these studies [41, 72, 73], here, we found reduced modulation of emotion judgment RT in patients with depression (**Fig. 6j**), which further inferred a reduced amygdala-dmPFC functional connectivity (**Fig. 7b**).

Increased connectivity between the amygdala, especially the basolateral amygdala, and distributed brain systems (including the PFC) involved in attention, emotion perception, and regulation, is associated with high childhood anxiety [74]. Individuals with anxiety show aberrant coupling between the amygdala and dmPFC during the presentation of images known to elicit negative affect [75]. Patients with generalized anxiety disorders show relatively lower intrinsic connectivity between the right amygdala and right PFC (both vmPFC and dmPFC) compared to controls [76]. However, in this study, we only observed a marginally significant modulation of emotion judgment RT (**Fig 6j**), which was not associated with a significant change in amygdala-PFC connectivity (**Fig. 7b**), likely due to the task that we used or the heterogeneity of patients that we sampled in the present study.

In patients with schizophrenia, resting-state fMRI has shown enhanced variability of intrinsic connectivity between the amygdala and vmPFC, which positively correlates with symptom severity [77]. On the other hand, task-based fMRI studies have demonstrated significantly weaker amygdala-PFC cortical coupling when processing negative distractors [78]. In this study, in both in-patient and out-patient SCZ patients, we observed a strongly reduced sensitivity to emotion intensity (**Fig. 6c**) as well as modulation of confidence rating (**Fig. 6e**), which may be attributed to enhanced amygdala-PFC functional connectivity (**Fig. 7a**).

Studies have shown abnormal amygdala-PFC connectivity in ASD [79, 80] and ADHD [38]. Both the PFC and amygdala are critical components of the “social brain” [81] and both brain regions be pathological in autism [82]. In humans, connections between these brain regions have been linked to reduced habituation after repeated presentations of faces in children with ASD [37]. Furthermore, children with ASD show reduced amygdala-PFC functional connectivity when viewing emotional faces [79] and when at rest (see [83] for a review), as well as abnormal structural connections [84]. In children with ADHD, the behavioral deficits in emotion regulation were found to be associated with altered amygdala-vmPFC intrinsic functional connectivity [38]. A theoretical account is that the amygdala orchestrates cognitive processes based on social stimuli, but it requires information conveyed from the PFC about the context in which those stimuli occur. In the absence of such contextual input, the amygdala may inappropriately interpret social stimuli [85]. Therefore, abnormal connections between the amygdala and PFC may underlie social deficits that cascade beyond facial processing to include processing of other socially relevant stimuli. In this study, we observed reduced sensitivity to emotion intensity in participants with ASD (**Fig. 6c**) and ADHD (**Fig. 6i**), and we also observed modulation of confidence rating in participants with ASD (**Fig. 6e**). For both groups, we did not observe an altered predicted amygdala-PFC functional connectivity (**Fig. 7a, b**).

### Advantages of a unique combination of multimodal approaches

For fMRI data, two different but complementary methods (PPI and DCM) were used to assess the functional connectivity among the amygdala, dmPFC, and vmPFC when processing emotion ambiguity. PPI is an anatomically unconstrained (whole-brain), data-driven approach that does not provide directionality of any changes in connectivity between regions [53, 86]. DCM is an alternative method for analyzing PPI within hypothesis-driven models that overcome this limitation [87].

In addition to fMRI, the unique combination of single-neuron recordings and scalp EEG shows promise in characterizing the brain network at a finer time scale. Among various EEG connectivity measures, cross-channel coherence evaluates the functional synchronization of cortical connections [54], while cross-frequency coupling (CFC) reflects the coordination of low and high frequency brain rhythms that are entrained by both external events and internal cognitive processes [56, 88]. The theoretical importance of these two measures may reflect how one brain region or rhythm is talking to another and is further modulated by experimental conditions. EEG coherence provides an important estimate of functional interactions between neural systems in a frequency-selective manner [54]. It can yield information about network formation and functional integration across brain regions. The CFC provides a plausible mechanism for the long-range communication between fast, spike-based computation with slower external events and internal states, thus guiding perception, cognition, and action [56, 89–92]. Notably, the scalp-based connectivity analysis can be extended to a source level, which may convey new information about the origins of signals and signal flow that is comparable with fMRI-based measures.

In particular, single-neuron recordings have a significantly higher spatial and temporal resolution compared to previous human studies solely using neuroimaging techniques, and this approach has provided a key missing link between animal neurophysiology and human neuroimaging. Simultaneous recordings from multiple brain regions permitted latency analysis, which may explain our findings at the circuit level and help distinguish between stimulus-driven vs. goal-driven modulating processes, making possible the isolation of specific neural processes especially relevant to behavior. It is worth noting that the limitations of singleunit recordings have been complemented by fMRI and EEG: the limited spatial coverage were complemented by fMRI whole-brain analysis, and the small number of patients from whom single-neuron recordings were made was balanced by the large number of in-lab EEG participants. Future studies are needed to perform detailed functional connectivity analysis with single-neuron data.

Together, the present study employed multimodal approaches that well complemented each other to comprehensively study the neural mechanisms of emotion ambiguity. It provided a systematic understanding of the amygdala-PFC network underlying emotion ambiguity with fMRI-based connectivity, EEG coordination of cortical regions, synchronization of brain rhythms, directed information flow of the source signals, and latency of single-neuron responses.

### Possible caveats

With the advantages of our study in mind, we would also like to note several limitations and possible caveats of our study. First, although we employed multimodal approaches to study functional connectivity that pointed to coherent results, they were not employed in the same participants. A future study is needed to account for the individual differences from different participant groups. Second, the online participants with neuropsychiatric disorders were only self-identified, and they may have large variance in their symptoms and severity. This may explain the weaker findings in these participants compared to the in-lab group. A future study is needed to confirm our findings with formally diagnosed patient groups. However, online participants are more representative of the general population so they may have a better generalizability to the general population. Lastly, we only predicted functional connectivity in patients with neuropsychiatric disorders based on functional connectivity from controls and patients’ behavior. A future study is needed to directly measure functional connectivity in these patients when they perform the same task.

## Author Contributions

S.S., R.Y., and S.W. designed experiments. S.S. and S.W. performed research. S.S., H.Y., and S.W. analyzed data. S.S., H.Y., R.Y., and S.W. wrote the paper. All authors discussed the results and contributed toward the manuscript.

## Acknowledgements

We thank Runnan Cao for help with online data collection and Ueli Rutishauser for providing the human single-neuron data. This research was supported by the AFOSR (FA9550-21-1-0088), NSF (BCS-1945230, IIS-2114644), NIH (R01MH129426), and Dana Foundation (to S.W.), and FRIS Creative Interdisciplinary Collaboration Program (No. 56045092), Operational Budget of President’s Discretionary Funds at Tohoku University (No. 56045090) and Japan Society for the Promotion of Science Grant-in-Aid for Early-Career Scientists (No. 22K15626) (to S.S.). The funders had no role in study design, data collection and analysis, decision to publish, or preparation of the manuscript.

## Supplementary Figures

**Supplementary Fig. 1.** Psychophysiological interaction (PPI) analysis for emotion (i.e., fear) intensity. **(a)** Decreasing fear intensity was correlated with increasing BOLD activity in the left dorsomedial prefrontal cortex (dmPFC), left anterior insular, and left amygdala. **(b)** PPI analysis revealed functional connectivity between the amygdala and dmPFC (peak: *x* = –9, *y* = 24, *z* = 48; 13 voxels, SVC, FWE P = 0.05).

**Supplementary Fig. 2.** A list of dynamic causal modeling (DCM) models that were analyzed in the present study.

**Supplementary Fig. 3**. Control DCM analysis with the right dorsomedial prefrontal cortex (dmPFC) and ventromedial prefrontal cortex (vmPFC). Legend conventions as in **Fig. 2**.

**Supplementary Fig. 4.** Reaction time (RT) and confidence rating (CR) for each condition and each participant group. Error bars denote ±SEM across participants. Shown above each plot are the p-values from one-way repeated measures ANOVA across conditions.

