## Supplementary figures and images for "Functional connectivity between the amygdala and prefrontal cortex underlies processing of emotion ambiguity"

### Supplemental Figure 1

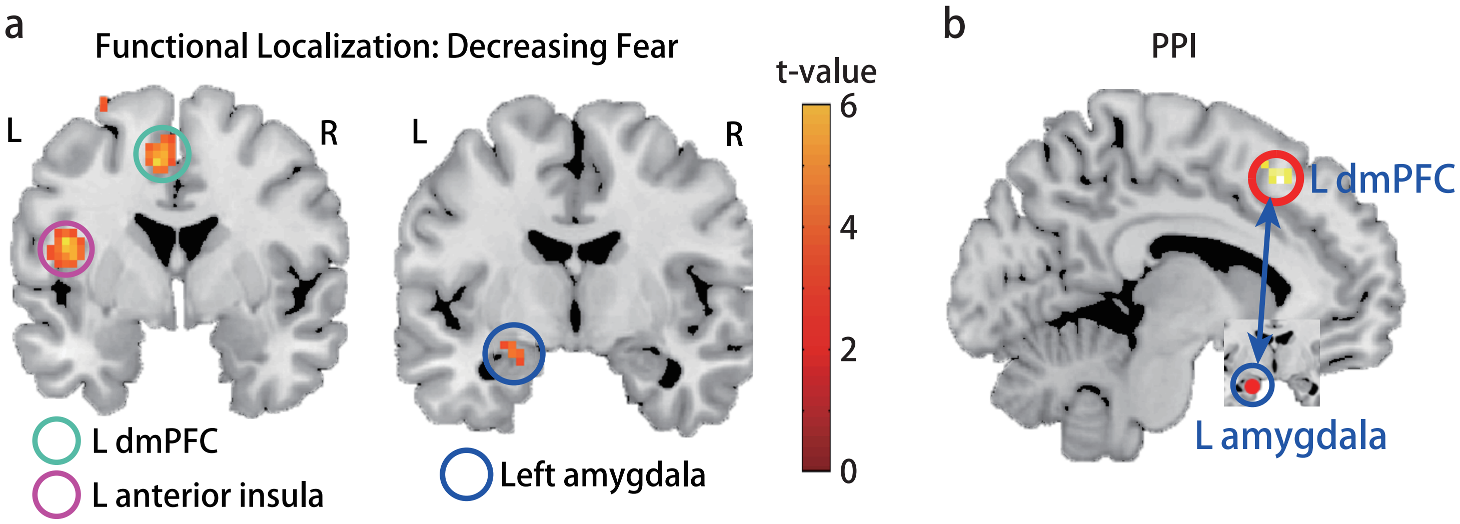

### Supplemental Figure 2

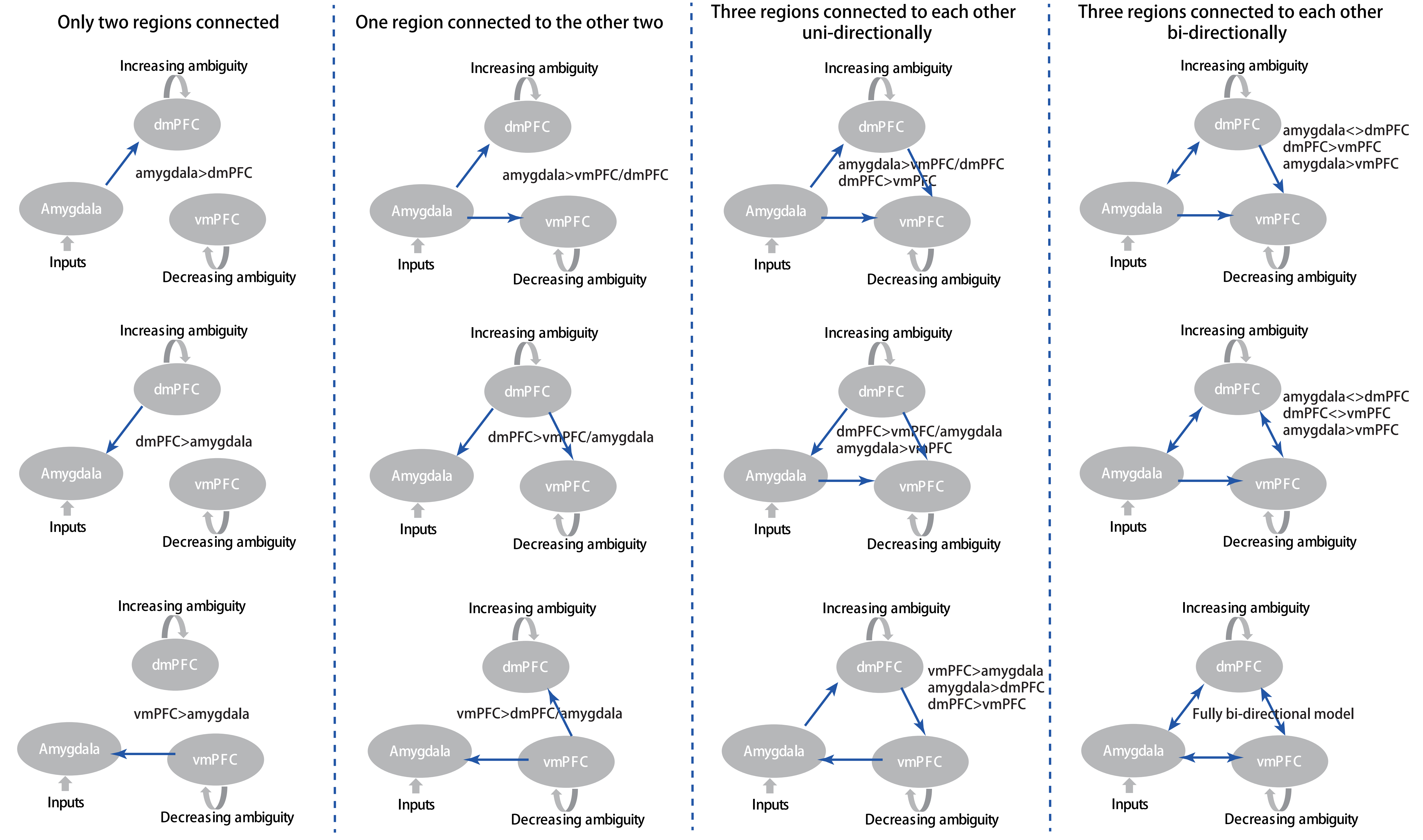

### Supplemental Figure 3

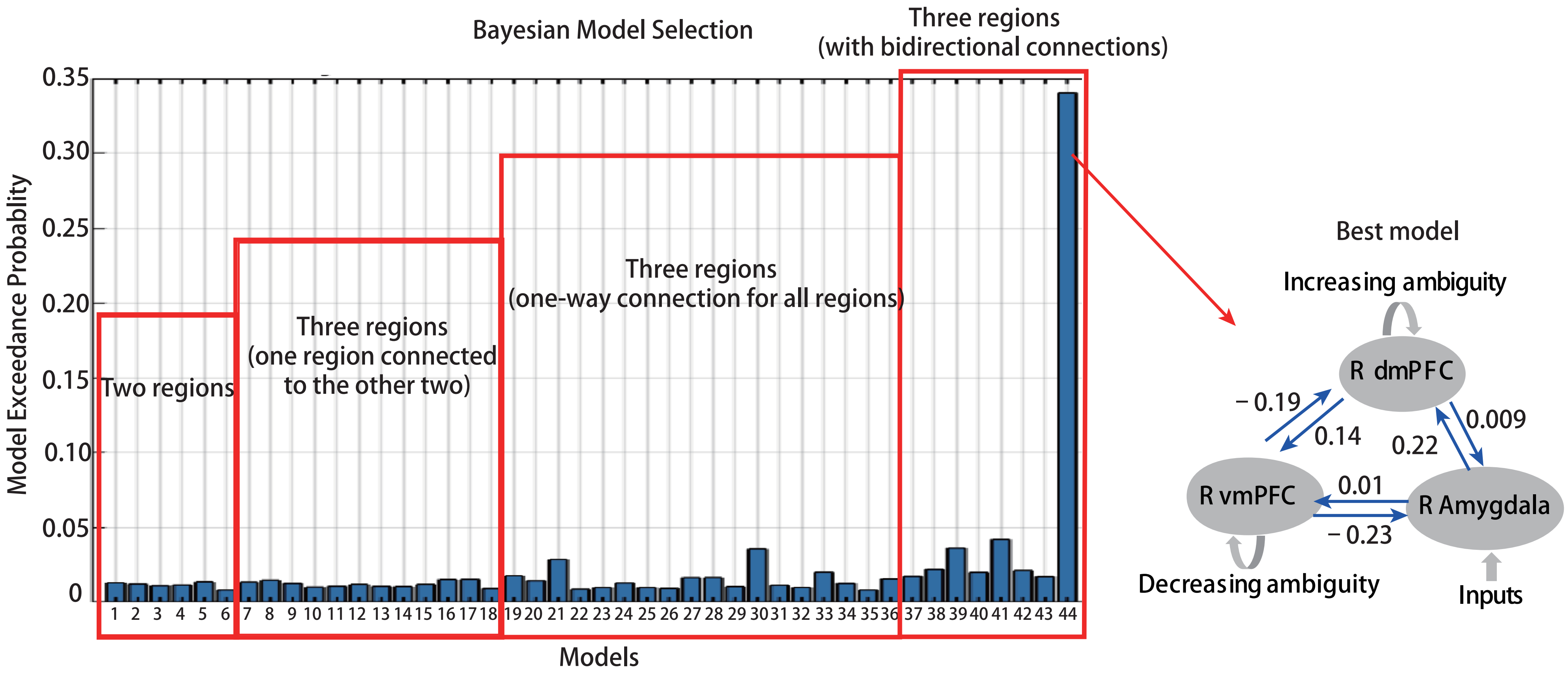

### Supplemental Figure 4

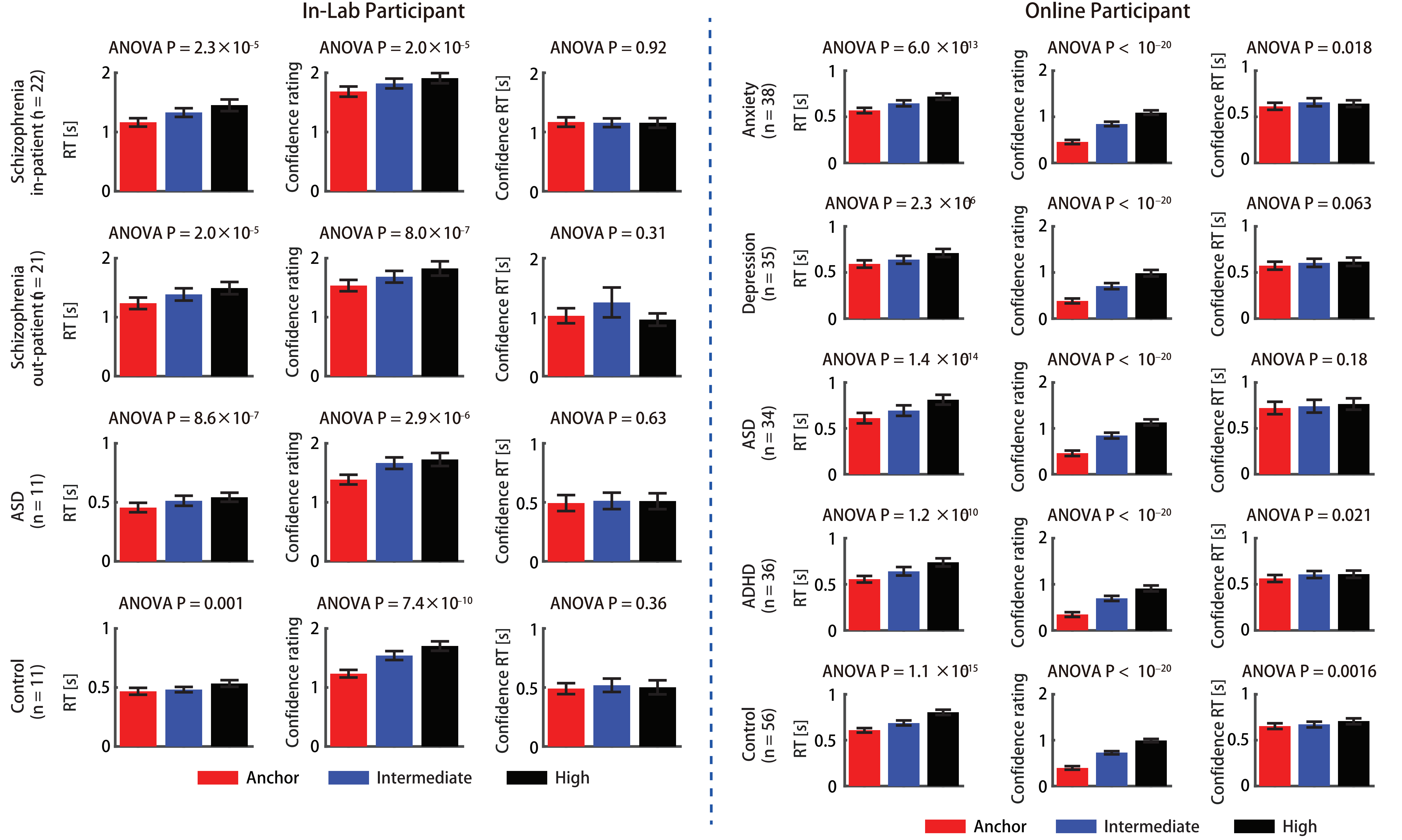
